# Structure and dynamics of FCHo2 docking on membranes

**DOI:** 10.1101/2021.04.20.440640

**Authors:** F. El Alaoui, I. Casuso, D. Sanchez-Fuentes, C. André-Arpin, R. Rathar, V. Baecker, A. Castro, T. Lorca, J. Viaud, S. Vassilopoulos, A. Carretero-Genevrier, L. Picas.

## Abstract

Clathrin-mediated endocytosis (CME) is a central trafficking pathway in eukaryotic cells regulated by phosphoinositides. The plasma membrane phosphatidylinositol-4,5-bisphosphate (PI(4,5)P_2_) plays an instrumental role in driving CME initiation. The F-BAR domain only protein 1 and 2 complex (FCHo1/2) is among the early proteins that reach the plasma membrane, but the exact mechanisms triggering its recruitment remains elusive. Here, we show the molecular dynamics of FCHo2 self-assembly on membranes by combining bottom-up synthetic approaches on *in vitro* and cellular membranes. Our results indicate that PI(4,5)P_2_ domains assist FCHo2 docking at specific membrane regions, where it self-assembles into ring-like shape protein patches. We show that binding of FCHo2 on cellular membranes promotes PI(4,5)P_2_ clustering at the boundary of cargo receptors and that this accumulation enhances clathrin assembly. Thus, our results provide a mechanistic framework that could explain the recruitment of early PI(4,5)P_2_-interacting proteins at endocytic sites.

## Introduction

The biogenesis of clathrin-coated vesicles requires precise and coordinated recruitment of more than ~ 50 different proteins to undergo the bending, elongation, and fission of the plasma membrane(1, 2). Different factors assist in recruiting endocytic proteins, such as the interaction with phosphoinositides(3), curvature sensing, and protein-protein interactions(4). Also, the membrane tension that results from membrane-cytoskeleton adhesion sets the load of forming transport vesicles(5). Although the initiation of endocytosis is a critical step, the exact mechanism triggering the nucleation of endocytic proteins at the plasma membrane is not well understood. The early stages of CME entail the nucleation of adaptor and accessory proteins, cargo, and lipids to undergo the bending of the plasma membrane(6). Several studies have shown that FCHo1/2 (Fer/CIP4 homology domain only protein 1 or 2) is among the early proteins recruited at endocytic sites(1, 7), where it establishes a network of interactions with pioneer proteins, such as Eps15, adaptor protein 2 (AP2), and transmembrane cargo(1, 8, 9). FCHo paralogs associate with membranes through the dimerization of F-BAR domains displaying a shallow concave surface that interacts with acidic phospholipids(7, 10, 11). A polybasic motif follows the F-BAR scaffold and provides a selective recognition for PI(4,5)P_2_(8). Finally, FCHo1/2 is flanked at the C-terminal by a μ-homology domain (μ-HD) that directly binds with multiple early endocytic proteins such as Eps15, intersectin 1 or CALM(7, 8, 12). Indeed, FCHo1/2 is required to recruit Eps15 on membranes(13), and the assembly of a FCHo1/2-Eps15-AP2 complex is essential to drive efficient cargo loading(8). The recruitment of FCHo1/2 on membranes is central to initiate the endocytic activity but, the underlying molecular mechanism remains unclear.

Here we combined sub-diffraction microscopy and high-speed atomic force microscopy (HS-AFM) with *in vitro* and *in cellulo* systems to show that PI(4,5)P_2_ domains regulate FCHo2 docking on flat membranes, where it self-assembles into ring-like shaped protein structures that are compatible with the size and temporal scale of clathrin-mediated endocytosis. Our results indicate that, in the absence of metabolizing enzymes, FCHo2 can engage a local PI(4,5)P_2_ enrichment at the boundaries of clathrin-regulated cargo receptors and enhance the formation of clathrin-positive assemblies. Finally, manipulation of membrane curvature through lithographic approaches showed that PI(4,5)P_2_ promotes the partition of FCHo2 at the edges of dome-like structures. Collectively, our work points out PI(4,5)P_2_ lateral lipid heterogeneities as an organizing mechanism supporting the docking and self-organization of PI(4,5)P_2_-interacting proteins that, like FCHo2, participate in the initial stages of CME.

## Results

### PI(4,5)P_2_ domains drive FCHo2 docking on membranes

To study FCHo1/2 recruitment on cellular membranes, we monitored by airyscan microscopy the binding of recombinant full-length FCHo2-Alexa647 on plasma membrane sheets. We generated plasma membrane sheets by ultrasound-mediated unroofing of cells stably expressing the transferrin receptor (TfR-GFP) as a model of cargo receptor regulated by FCHo1/2(7). To monitor the subcellular dynamics of lipids we used TopFluor fatty acid conjugates, as previously reported(14). We loaded HT1080 unroofed cells with TF-TMR-PI(4,5)P_2_ or fluorescent phosphatidylserine as a control (PS, TF-TMR-PS) (Figure 1A and B). The functionality of recombinant FCHo2 was determined by performing a tubulation assay using membrane sheets made of brain polar lipids, as previously reported(15) (Figure S1). Kymograph analysis on plasma membrane sheets showed that FCHo2 is preferentially recruited to PI(4,5)P_2_-enriched regions often co-localized with the TfR (Figure 1C). We found that on these regions the kinetics of FCHo2 recruitment was faster than in the absence of PI(4,5)P_2_ enrichment (Figure 1D). Indeed, the appearance of FCHo2 puncta on plasma membrane sheets showed a positive correlation with the intensity of PI(4,5)P_2_ compared to PS (Figure 1E).

**Figure 1.**
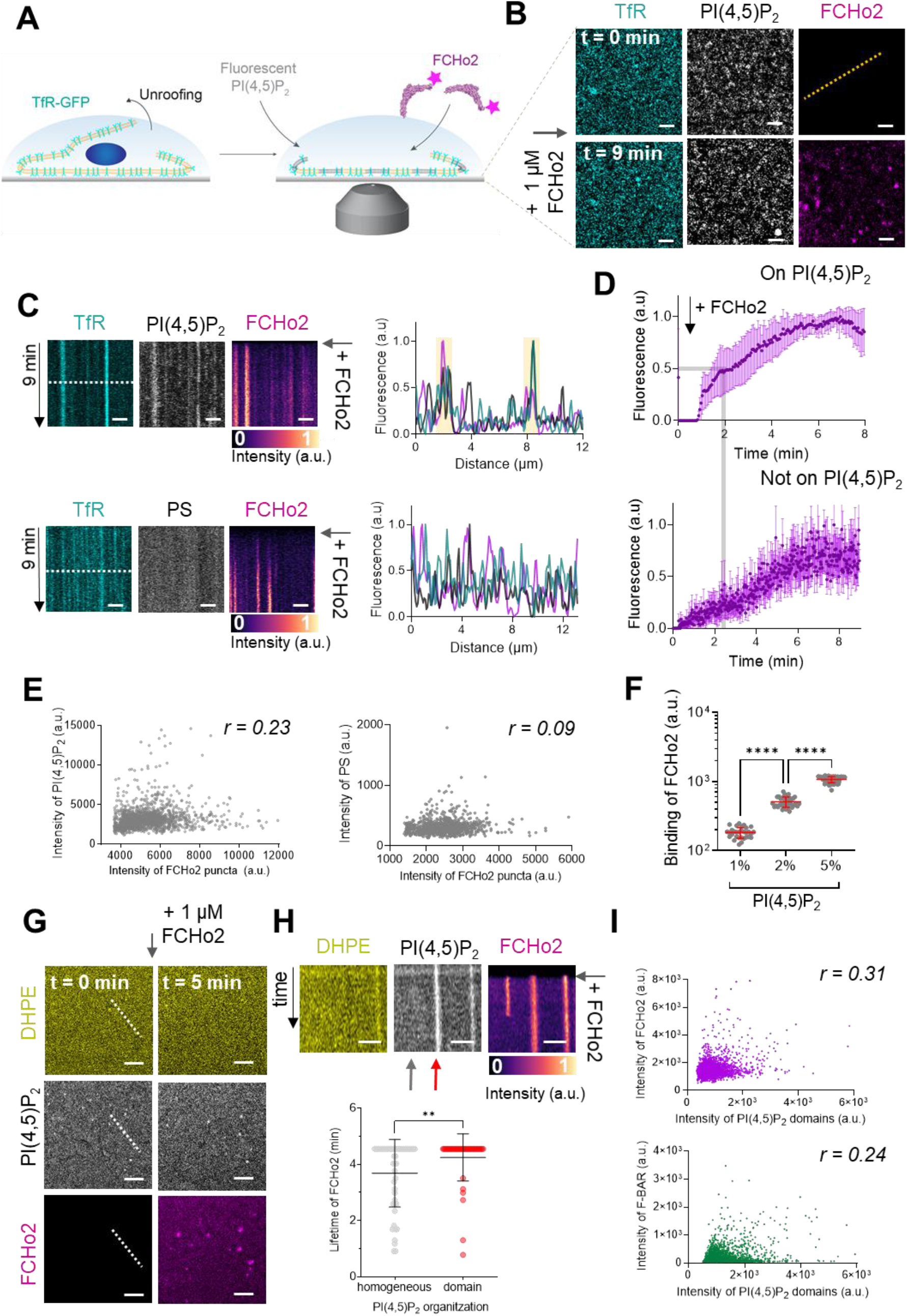
FCHo2 is recruited to PI(4,5)P_2_-enriched regions on cellular and *in vitro* membranes. (A) Cartoon of the experimental setup to monitor the recruitment of FCHo2-Alexa647 on cellular membranes doped with fluorescent PI(4,5)P_2_. (B) Representative airyscan time-lapse showing the dynamics of PI(4,5)P_2_ (gray), FCHo2 (magenta) and TfR-GFP (cyan) on plasma membrane sheets. Scale bar, 5 μm. (C) *Top*, kymograph analysis along the dashed line in B on plasma membrane sheets doped with TF-TMR-PI(4,5)P_2_ and intensity profile of the TfR (cyan), PI(4,5)P_2_ (gray) and FCHo2 (magenta) along the dashed line in the kymograph. Time scale is 9 min. *Bottom*. Representative kymograph analysis on plasma membrane sheets doped with TF-TMR-PS and intensity profile of the TfR (cyan), PS (gray) and FCHo2 (magenta) along the dashed line in the kymograph. Time scale is 9 min. Scale bar, 2 μm. (D) Fluorescence quantification over time of the FCHo2 recruitment (magenta) on PI(4,5)P_2_-enriched regions (*top*) and not on PI(4,5)P_2_ (bottom) on plasma membrane sheets. Each curve represents the mean ± s.d., n = 7. (E) Scatter plots of the correlation between the intensity of FCHo2 puncta and PI(4,5)P_2_ or PS intensity. n=11448 and n=1090, respectively. The correlation coefficient was r = 0.23 and r = 0.09, respectively. (F) Binding of FCHo2 on lipid membranes containing different % mol of PI(4,5)P_2_ (20% of total negative charge). Mean ± s.d. in red. one-way ANOVA (****, P < 0.0001). n=31, n=39, and n=54 for 1%, 2%, and 5% mol of PI(4,5)P_2_, respectively. (G) Airyscan time-lapse showing the dynamics of FCHo2 (magenta) on 5% of PI(4,5)P_2_ membranes doped with fluorescent PI(4,5)P_2_ (gray) and DHPE (yellow). Scale bar, 2 μm. (H) *Top,* kymograph analysis along the white dashed line in F. Time scale, 5 min. Scale, 2 μm. *Bottom,* lifetime of FCHo2 (in min) relative to the PI(4,5)P_2_ organization: homogeneous (grey) and on domains (red), as highlighted by the arrows in the corresponding kymograph image. Mean ± s.d. in black. Welch’s t-test (**, P = 0.0098). n=50, n=43 for a homogeneous and on domains, respectively. (I) Scatter plots of the correlation between the intensity of FCHo2 (purple) and F-BAR (green) relative to the intensity of PI(4,5)P_2_ domains on 5% mol PI(4,5)P_2_-membranes. n=11448 for FCHo2 and n=12828 for F-BAR. The correlation coefficient, r, was r = 0.31 and r = 0.24, respectively.

To determine if the spatial recruitment of FCHo2 is mediated by PI(4,5)P_2_-enriched domains, we first estimated the protein binding on supported lipid bilayers made of 20% of total negatively charged lipids and different % mol of PI(4,5)P_2_. As expected, increasing amounts of PI(4,5)P_2_ favored FCHo2 binding on flat membranes (Figure 1F). Next, we analyzed the spatial recruitment of FCHo2 on 5% PI(4,5)P_2_-containing supported lipid bilayers doped with 0.2% mol of fluorescent PI(4,5)P_2_ (TF-TMR-PI(4,5)P_2_) and 0.2% of Oregon Green 488 1,2-Dihexadecanoyl-sn-Glycero-3-Phosphoethanolamine (OG-DHPE) at the expenses of their non-labeled counterpart (Figure S2). As a result, we designed as PI(4,5)P_2_-enriched domains the existence of spots that displayed a local increase in the TF-TMR-PI(4,5)P_2_ signal along with a decrease in the intensity of OG-DHPE, as compared to a homogeneous distribution of lipids characterized by an equivalent partitioning of both lipid dyes (Figure S2). We characterized the steady-state organization of the fluorescent PI(4,5)P_2_ into small domains as compared to other anionic lipids (Figure S3), in agreement with previous numerical simulations(16). The temporal analysis of FCHo2 recruitment (Figure 1G) showed that binding to PI(4,5)P_2_ domains resulted in the formation of long-lived FCHo2 puncta (red arrow, Figure 1H), whereas its association with a homogenous distribution of PI(4,5)P_2_ was more likely to lead to FCHo2 disassembly (gray arrow, Figure 1H).

Next, we analyzed the contribution of the F-BAR domain on the preferential localization of the full-length protein on PI(4,5)P_2_-domains (Figure 1I). Our results show that the binding of both proteins on membranes follows a positive correlation with the existence of PI(4,5)P_2_ domains. Although this correlation was more significant for FCHo2, the preferential recruitment of the F-BAR domain to PI(4,5)P_2_ enriched regions is in agreement with the enhanced binding of the domain at 5 % mol of PI(4,5)P_2_ (Figure S4). Collectively, these results indicate that PI(4,5)P_2_-enriched regions act as docking sites for FCHo2 recruitment on membranes through the electrostatic interaction of the F-BAR and polybasic motif.

### FCHo2-mediated PI(4,5)P_2_ clustering primes pre-endocytic events

The formation of PI(4,5)P_2_ clusters at the plasma membrane orchestrates the recruitment of PI(4,5)P_2_-binding proteins *via* ionic-lipid protein interactions(17, 18). Several structural domains of endocytic proteins, including the F-BAR domain of Syp1, locally accumulate PI(4,5)P_2_ on *in vitro* membranes(19, 20), and we showed that BIN1 recruits its downstream partner dynamin through this mechanism. We thus investigated the impact of FCHo2 on PI(4,5)P_2_ clustering formation. Injection of FCHo2-Alexa647 on TfR-GFP plasma membrane sheets led to the binding of the protein and the formation of sub-micrometric FCHo2-positive puncta that co-localized with PI(4,5)P_2_ and the cargo receptor (Figure 2A). Analysis of FCHo2 dynamics showed that the protein binding convoyed an increase of the PI(4,5)P_2_ signal on TfR-GFP-positive puncta but not on PS labeled plasma membrane sheets. To determine if FCHo2-mediated PI(4,5)P_2_ enrichment was a general feature of FCHo2, we monitored its binding relative another clathrin-regulated cargo, the EGF receptor (EGFR) (Figure 2B). In this case, we observed a concomitant increase of both the PI(4,5)P_2_ and EGFR signal at FCHo2-postivie puncta (Figure 2B and S5), possibly as a result of the interaction of the EGFR juxta-membrane domain with PI(4,5)P_2_(21). Quantification of the intensity of PI(4,5)P_2_ domains confirmed a local increase in the PI(4,5)P_2_ signal after the addition of FCHo2 on cellular membranes (Figure 2C). Therefore, pointing out that membrane binding of FCHo2 induces P(4,5)P_2_ clustering at the boundary of clathrin-regulated cargo receptors(11, 22).

**Figure 2.**
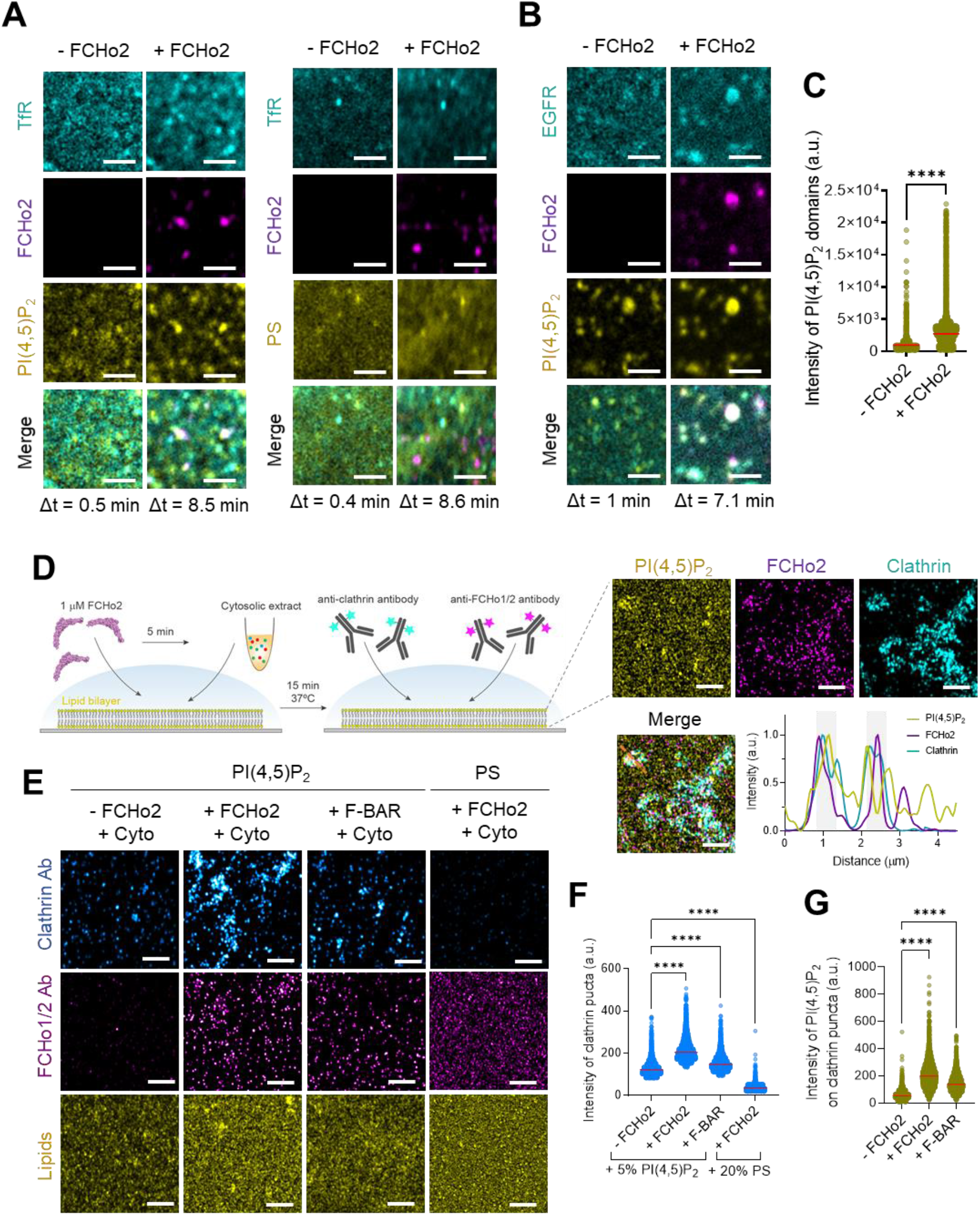
FCHo2 induces PI(4,5)P_2_ clustering on cargo receptors and clathrin-positive puncta. (A, B) Representative z-projected time-lapse confocal images of plasma membrane sheets showing the co-localization of PI(4,5)P_2_ or PS (yellow) and clathrin-regulated cargoes TfR or EGFR (cyan) before (- FCHo2) and after the addition of FCHo2 (+ FCHo2, magenta). The projected time interval (Δt) is specified below each image. Sale bar, 2 μm. (C) Distribution of the intensity of PI(4,5)P_2_ domains before (-FCHo2) and after addition of FCHo2 (+ FCHo2) on plasma membrane sheets. Mean ± s.d, in red. Welch’s t-test (****, P < 0.0001). n=4397 and n=33815, respectively. (D) Cartoon of the *in vitro* clathrin-coat assembly with cytosolic extracts and immunofluorescence assay to detect FCHo2 and clathrin-positive structures on 5% mol PI(4,5)P_2_-containing supported lipid bilayers. Representative airyscan images on lipid bilayers doped with fluorescent PI(4,5)P_2_ (yellow) showing the co-localization with FCHo2 (magenta) and clathrin (cyan). Intensity profile along the orange dotted line in the corresponding image. Sale bar, 2 μm. (E) Representative airyscan images of lipid bilayers containing either 5% mol PI(4,5)P_2_ or 20% PS (yellow) (total negative charge is 20%) that were incubated with recombinant FCHo2 (+ FCHo2) or F-BAR domain (+ F-BAR) or without FCHo2 (- FCHo2) before addition of 4 mg/ml of cytosolic extracts (+ Cyto). Clathrin (blue) and FCHo1/2 (magenta) were detected by immunofluorescence (antibody, Ab). Sale bar, 2 μm. (F) Distribution of the intensity of clathrin structures in the presence or absence of FCHo2 or F-BAR on PI(4,5)P_2_ or PS-containing bilayers. Mean ± s.d. is displayed in red and one-way ANOVA in black (****, P < 0.0001). n=7784, n =7539, n=4798 and n=6642, as in the graph. (G) Distribution of the intensity of fluorescent PI(4,5)P_2_ on clathrin structures in the presence or absence of FCHo2 or F-BAR. Mean ± s.d., in red. One-way ANOVA (****, P < 0.0001). n=7784, n =7539 and n=6642, as in the graph.

We next analyzed if FCHo2-mediated PI(4,5)P_2_ clustering participates in the formation of clathrin-coated structures. To this end, we reconstituted *in vitro* clathrin-coat assembly by using cytosolic extracts, as previously reported(23, 24). We supplemented active cytosolic components from *Xenopus* egg extracts with a freshly prepared “energy mix”, as detailed in the methods section. We determined by immunofluorescence the formation of clathrin-positive puncta on supported lipid bilayers containing 5% mol PI(4,5)P_2_ (including 0.2% of fluorescent PI(4,5)P_2_) previously incubated with or without non-labeled FCHo2 (Figure 2D). As expected, in the absence of FCHo2 the addition of cytosolic extracts leads to the appearance of clathrin-positive puncta, as compared to the energy mix alone (Figure 2E and S6). Under these conditions, we could also detect a residual signal of the FCHo1/2 antibody, corresponding to the endogenous protein present in the cytosolic extracts. Incubation of 5% PI(4,5)P_2_-containing lipid bilayers with 1 μM of FCHo2 resulted in a 1.7-fold increase in the mean intensity of clathrin-positive spots (Fig. 2E and F), which convoyed by a 3.6-fold local increase in the PI(4,5)P_2_-signal. We could detect a 2.5-fold increase in the local PI(4,5)P_2_ intensity in the presence of the F-BAR domain alone, which also lead to a moderate increase in the appearance of clathrin-positive spots, in agreement with the ability of the F-BAR domain in PI(4,5)P_2_ clustering formation(19). Although FCHo2 was still able to bind on members, replacement of PI(4,5)P_2_ by PS prevented the detection of clathrin in our *in vitro* assay, supporting the functional role of PI(4,5)P_2_ in clathrin assembly on membranes. Thus, in addition to its membrane bending ability, these data indicate that FCHo2 might promote clathrin assembly by clustering PI(4,5)P_2_.

### Assembly of FCHo2 into molecular clusters prompts membrane bending

A characteristic hallmark of endocytic proteins on cellular membranes is their spatial organization into punctate structures(7, 8), and we systematically observed this feature on plasma membrane sheets and *in vitro* membranes (Figure 1 and 2). Co-segregation of early endocytic proteins into sub-micrometer scale clusters relies on multivalent interactions(13). Therefore, we asked what might be the role of PI(4,5)P_2_ in the spatial organization of FCHo2. We estimated by airyscan microscopy the average size of FCHo2 puncta on cellular membranes (i.e. plasma membrane sheets) and on 5% PI(4,5)P_2_-containing bilayers after the addition of 1 μM of FCHo2-Alexa 647 (Figure 3A and B), which was on both cases ~ 0.067 μm^2^. On lipid bilayers doped with 5% PI(4,5)P_2_ we could also detect that the F-BAR domain alone can assemble into punctate structures, although the average size was ~ 0.094 μm^2^. Interestingly, replacing 5% PI(4,5)P_2_ by PS lead to a homogeneous surface distribution of FCHo2 and prevented the detection of sub-micrometric puncta. Therefore, suggesting that PI(4,5)P_2_ multivalent interactions through the F-BAR promote FCHo2 segregation into molecular clusters.

**Figure 3.**
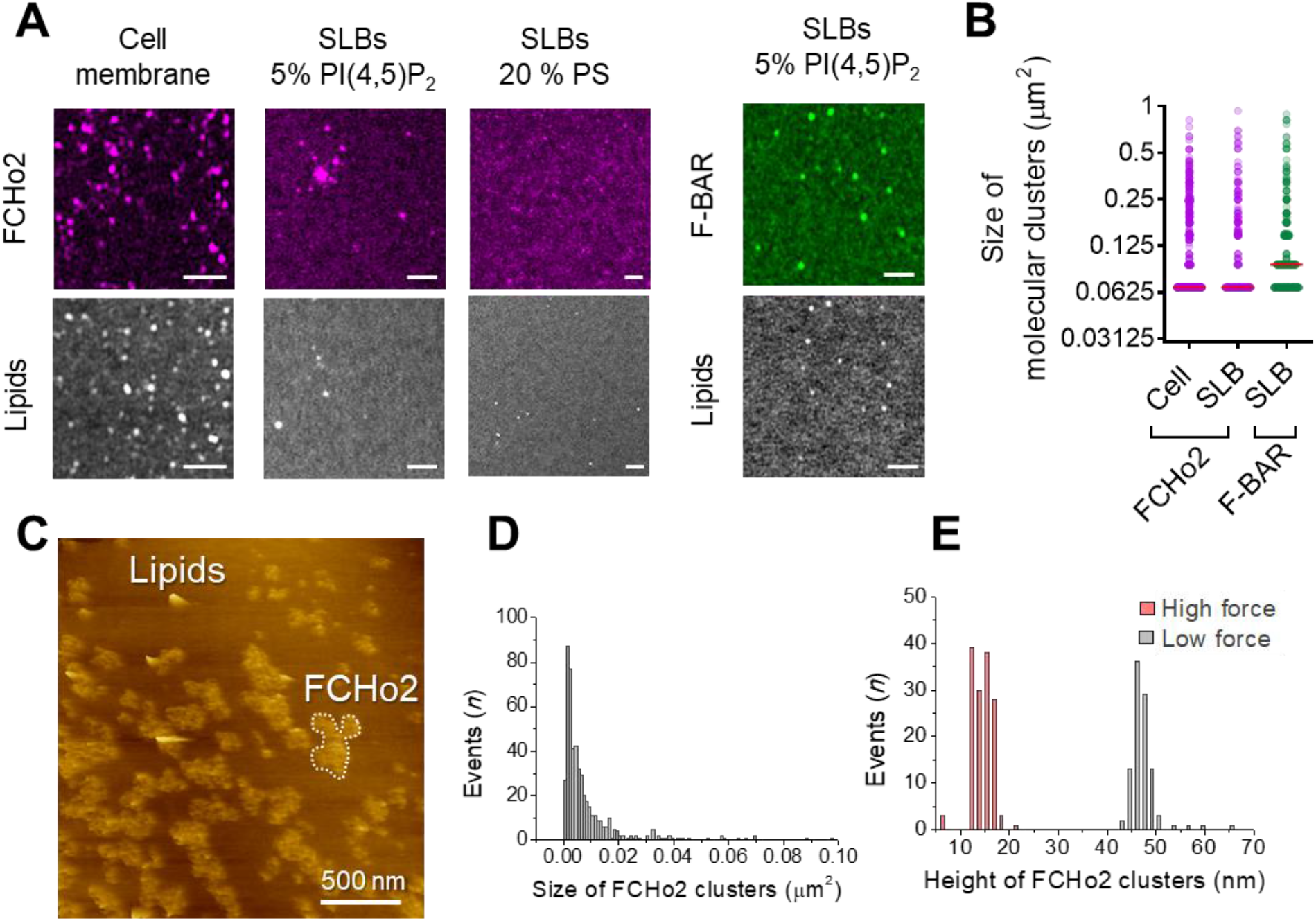
FCHo2 forms molecular clusters on PI(4,5)P_2_-containing bilayers. (A) Representative airyscan images of plasma membrane sheets or lipid bilayers doped with either 5% mol PI(4,5)P_2_ or 20% PS (total negative charge is 20%) and incubated with FCHo2-Alexa647 (magenta) or F-BAR-Alexa647 (green). Sale bar, 2 μm. (B) Size distribution of FCHo2 (magenta) and F-BAR (green) molecular clusters (in μm²) on plasma membrane sheets (cell) or lipid bilayers (SLB) containing 5 % mol of PI(4,5)P_2_. Median value is displayed in red. n=10793, n=19287, and n=7028, as in the graph. (C) Representative AFM image of FCHo2 molecular clusters (white dashed region) on supported lipid bilayers containing 5% mol PI(4,5)P_2_. (D) Size distribution of FCHo2 molecular clusters (in μm^2^) obtained from AFM images. (E) Height distribution of FCHo2 molecular clusters (in nm) at low (gray) and high (red) setpoint forces.

Next, we used atomic force microscopy (AFM) to investigate FCHo2 molecular clusters at nanometer-scale resolution. Supported lipid bilayers doped with 5% PI(4,5)P_2_ were formed on freshly cleaved mica disks (see methods section). Before adding proteins, we confirmed the homogeneity and absence of defects of supported bilayers under the imaging buffer. Injection of full-length FCHo2 at 1 μM in the imaging chamber resulted in sub-micrometric protein patches with a median dimension of ~ 0.005 μm^2^ that protruded out of the flat membrane surface with an average height of ~ 47 ± 3 nm (Figure 3C-E). An increase in the setpoint force from minimal values (a few tens of pN) to intense forces (around one hundred pN) resulted in the reduction of the height of FCHo2 clusters down to ~ 15 ± 2 nm, thus suggesting that FCHo2 can moderately bend supported lipid membranes at minimal AFM imaging forces.

### FCHo2 forms ring-like shape assemblies on flat membranes

To establish the biogenesis of FCHo2 clusters at the molecular level, we used high-speed AFM (HS-AFM). Real-time imaging of the initial stages revealed the entire molecular process of FCHo2 cluster formation, from the binding of single FCHo2 homodimers to the growth of molecular clusters (Figure 4A). Representative time-lapse images and kymograph analysis along the dashed region at t = 0 s showed that the binding of individual FCHo2 proteins engages an indentation of few nm in the lipid membrane adjacent to the protein surface (Figure 4B, green arrowheads), as delineated from the cross-section profile along the red dashed box in the corresponding kymograph (Figure 4A). The binding of FCHo2 rapidly prompted by the arrival of additional FCHo2 homodimers (as highlighted by white arrowheads in the kymograph). This stage of the process was characterized by minimal lateral interactions and the existence of contacts between adjacent FCHo2 homodimers, ultimately leading to a ring-like organization (Figure 4C). This dynamic reorganization that we named the “ring formation” step spanned over ~ 80-100 s.

**Figure 4.**
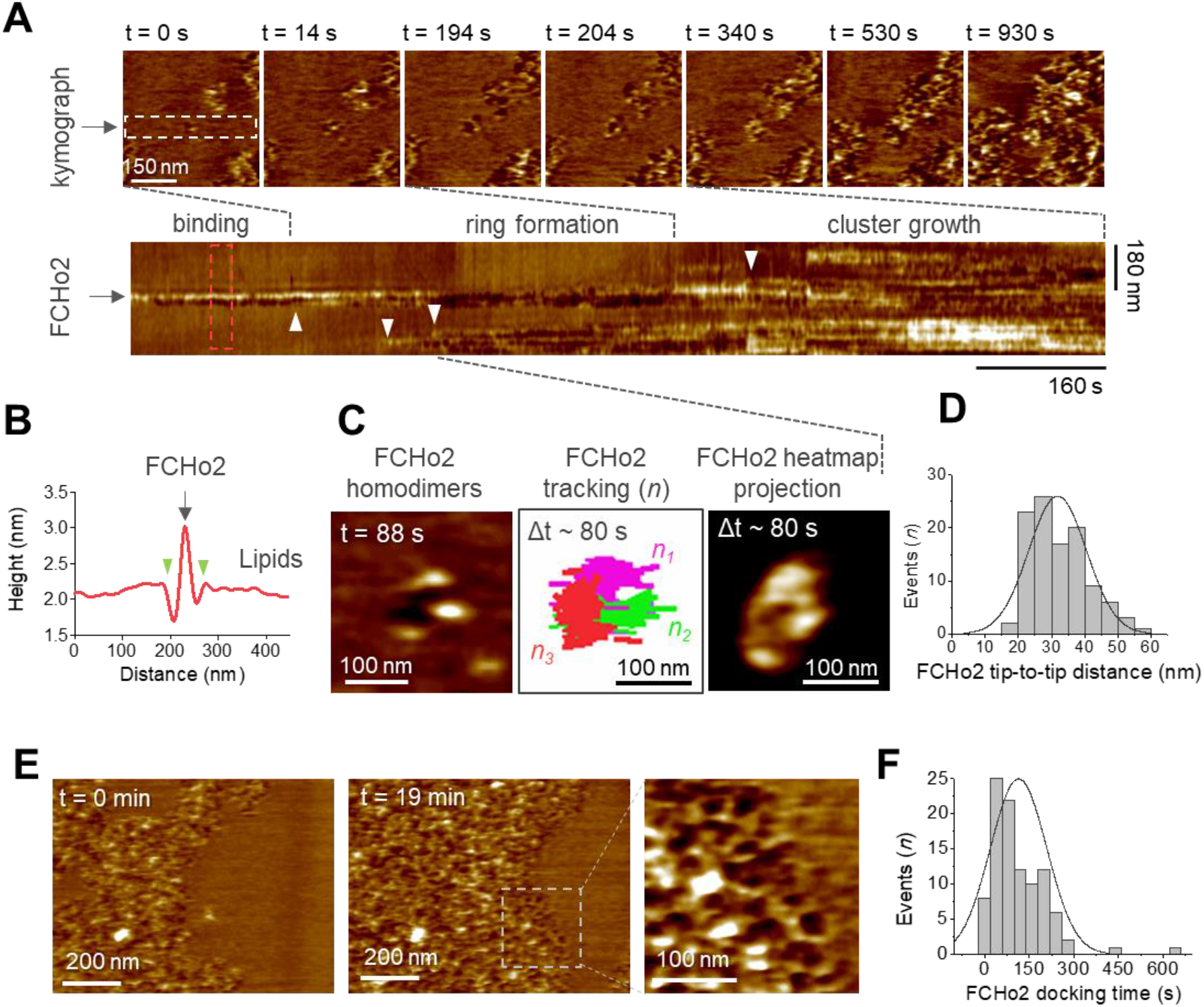
Molecular dynamics of FCHo2 self-assembly on PI(4,5)P_2_-containing membranes. (A) HS-AFM movie frames of the binding of FCHo2 on 5% mol PI(4,5)P_2_-containing membranes. Kymograph analysis performed on the white outline to display the representative stages of FCHo2 binding and self-assembly on flat membranes. White arrowheads highlight the docking of new FCHo2 homodimers to the growing molecular cluster. (B) Profile analysis along the red outline in A. Green arrowheads highlight the membrane invagination upon binding of FCHo2 on lipid membranes. (C) HS-AFM movie snapshot of the FCHo2 ring formation along with a representative tracking of individual homodimers (*n* = 3) and the heat map projection within a time interval, Δt, of ~ 80 s. (D) Size distribution (in nm) estimated from individual FCHo2 proteins before the cluster growth (t ≤ 200 s). (E) HS-AFM snapshots (t = 0 min and t = 19 min) illustrating the growth of FCHo2 molecular clusters into ring-like shape protein patches. Magnified image corresponds to the dashed outline. (F) Distribution of the docking time (in s) of individual and ring-like FCHo2 assemblies during the cluster growth.

The identification of individual proteins at the initial stages allowed us to extract the average dimension of the full-length protein interacting with flat membrane, which was ~ 32 ± 8 nm (Figure 4D) and in good agreement with the size of the F-BAR domain reported from electron microscopy micrographs (10). After the ring formation, we observed the growth of FCHo2 clusters through docking events that involved individual FCHo2 homodimers and the coalescence of adjacent FCHo2 rings (Figure 4A, white arrowheads). FCHo2 self-organization into hollow ring-like assemblies was particularly discernible at the growth front of FCHo2 molecular clusters (Figure 4E and magnified image). Although the entire formation of FCHo2 molecular clusters expanded over few tens of minutes, we found that the docking of individual proteins and rings to support the expansion of the cluster took place every ~ 115 ± 94 s (Figure 4F). Collectively, our results suggest that on flat membranes, FCHo2 exhibits an intrinsic ability to self-assemble into a ring-like shape molecular complex independently of the local protein density.

### PI(4,5)P_2_ assists FCHo2 partitioning on curved membranes

Because the transition from a flat surface to a dome-like invagination is a major step in the formation of clathrin-coated structures(2), we set out to monitor the organization of FCHo2 on curved membranes. To this end, we engineered arrays of SiO_2_ vertical nano-domes of radii *R* ~ 150 nm using soft nano-imprint lithography (soft-NIL) (Figure 5A), as previously reported(25). We functionalized SiO_2_ nano-patterned substrates with supported lipid bilayers containing 20% of negatively charged lipids. First, we determined by airyscan microscopy the surface organization of the F-BAR domain and FCHo2 labeled with Alexa647 on nano-domes in the presence of lipid bilayers containing 5% mol of PI(4,5)P_2_ (Figure 5B). The three-dimensional (3D) rendering of the protein signal relative to a reference marker to depict the nano-dome topography (DHPE lipid dye), showed that while the F-BAR domain is excluded from the nano-structure, the FCHo2 staining is well present at the base of the nano-dome (Figure 5B). We hypothesize that if the association of FCHo2 with PI(4,5)P_2_ is essential for its localization on curved membranes, disrupting this interaction would change its spatial organization. Indeed, this was the case and replacing PI(4,5)P_2_ by PS (20% mol PS) lead to complete distribution of FCHo2 all over the nano-dome surface (Figure 5B, yellow).

**Figure 5.**
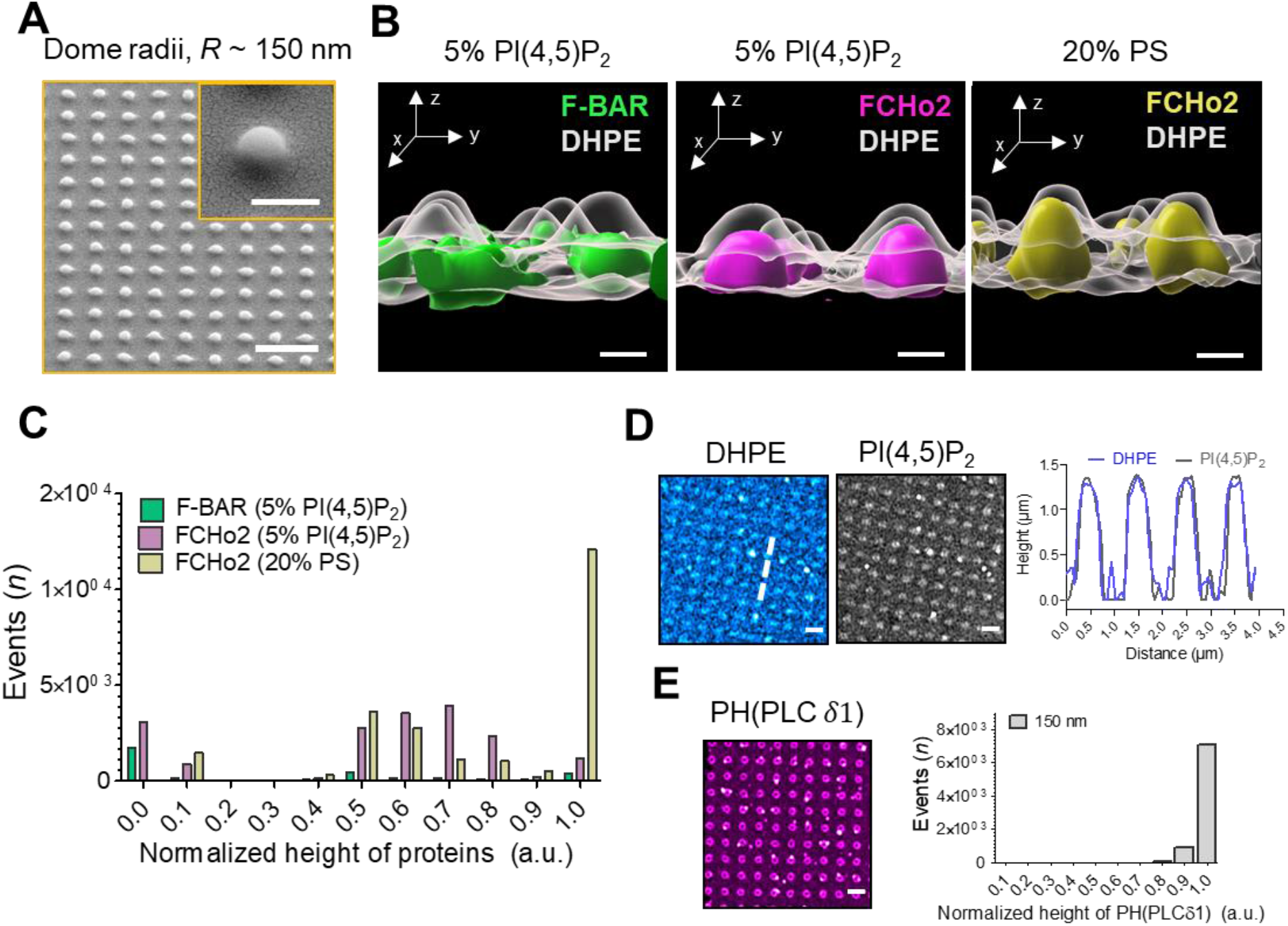
PI(4,5)P_2_ assists the organization of FCHo2 on curved membranes. (A) SEM images of SiO2 substrates displaying an array of nano-domes with a radius, *R* ~ 150 nm generated by soft-NIL. Scale bar, 2 μm. Inset, scale bar is 500 nm. (B) Representative 3D renders of the surface organization of the F-BAR domain (green) and FCHo2 (magenta) on nano-domes functionalized with either 5% mol PI(4,5)P_2_ or 20% PS-containing membranes (yellow) relative to the DHPE lipid dye signal (gray). Scale bar, 400 nm. (C) Distribution of the normalized maximal height of the surface distribution of the F-BAR domain (green) and FCHo2 (magenta) on nano-domes functionalized with 5% mol of PI(4,5)P_2_ or 20% PS lipid bilayers (yellow). n=3300, n=18145 and n=23043, respectively. (D) *Left,* z-stack intensity projected airyscan images of the distribution of fluorescent DHPE and PI(4,5)P_2_ on nano-domes. Scale bar, 2 μm. *Right,* cross-section of the normalized height of DHPE (blue) and PI(4,5)P_2_ (grey) lipid dyes obtained from the dashed line in the corresponding airyscan image. (E) *Left,* z-stack intensity projected airyscan images of the PH (PLCδ1) on nano-domes functionalized with 5% mol of PI(4,5)P_2_ lipid bilayers. Scale bar, 2 μm. *Right,* distribution of the normalized maximal height of the surface distribution of PH (PLCδ1) on nano-domes. n=7974.

To quantitatively estimate the preferential localization of FCHo2 on curved membranes, we computed for each nano-dome in patterned array the normalized maximal height of the protein signal relative to the DHPE lipid signal, which was used as a reference of the nano-dome height (see methods). Thus, a normalized maximal height close to ~ 1.0 indicates an increased probability of detecting the protein all over the nano-structure surface, whereas a value of ~ 0 would reflect an absence of preference for the nano-structure. On nano-domes functionalized with PI(4,5)P_2_-containing membranes, we obtained a median value of the F-BAR domain maximal height of ~ 0 (Figure 5C, green), whereas FCHo2 displayed two preferential localizations but with a median height on nano-domes that was of ~ 0.64 (Figure 5C, magenta). Finally, on membranes containing PS we obtained that the maximal median height of FCHo2 on nano-domes was ~ 1 (Figure 5C, yellow), in agreement with the 3D renderings in Figure 5B.

Next, we determined whether FCHo2 partitioning on nano-domes might be affected by a heterogeneous distribution of PI(4,5)P_2_ on curved membranes. To this end, we estimated the axial localization of the PI(4,5)P_2_ dye relative to the DHPE lipid dye on nano-domes coated with 5% PI(4,5)P_2_-containing lipid bilayer (Figure 5D). The cross-section analysis showed an equivalent surface distribution of the PI(4,5)P_2_ and DHPE lipid dye under our experimental conditions. To confirm that TF-TMR-PI(4,5)P_2_ signal was replicating the actual organization of the total pool of PI(4,5)P_2_ on nano-domes, we analyzed the distribution of the PH domain of PLCδ1 PH(PLCδ1), which is a well-established reporter of PI(4,5)P_2_(26). We obtained a median of the maximal height of the PH(PLCδ1) of ~ 1.0, indicating a homogenous distribution along the nano-dome surface (Figure 5F).

## Discussion

This study reports a molecular visualization of the docking and self-assembly of the endocytic protein FCHo2 on *in vitro* and cellular membranes (Figure 6). Our results show that PI(4,5)P_2_ is a primary spatial regulator of the recruitment of FCHo2 by promoting its accumulation and sorting on flat and curved membranes. These observations support the model that multivalent interactions play an instrumental role in upholding the early stages of endocytosis. We show that FCHo2 oligomerization induces PI(4,5)P_2_ clustering formation on cellular membranes that often co-localized with TfR and EGFR-positive puncta. This association was particularly remarkable in the case of the EGFR and agreed with the observation that electrostatic interaction of the polybasic motifs at the cytoplasmic tail of the EGFR mediates its clustering on PI(4,5)P_2_-enriched domains(16, 21, 27). FCHo1/2 is needed to recruit Eps15(7) and form FCHo1/2-Eps15 micrometer-scale domains on membranes(13). By driving PI(4,5)P_2_ clustering formation at the boundary of cargo receptors, FCHo2 is likely to improve the formation of a network of PI(4,5)P_2_-interacting proteins. Indeed, our observations showing that FCHo2 intensifies the formation of clathrin-positive assemblies on in vitro membranes through PI(4,5)P_2_-rich interfaces accords with previous studies pointing that clustering of Fcho1/2Eps15/AP_2_ primes endocytosis(8). We discerned clathrin-positive puncta in lower FCHo2 concentrations but not in PI(4,5)P_2_-depleted membranes, which agrees with the observation that AP_2_ can create its local pool of PI(4,5)P_2_(28), although it requires PI(4,5)P_2_ for its localization and activation at the plasma membrane(29, 30). Our measurements point that local PI(4,5)P_2_ enrichment induced by FCHo2 might operate as a complementary mechanism to enhance PI(4,5)P_2_-mediated interactions at endocytic sites.

**Figure 6.**
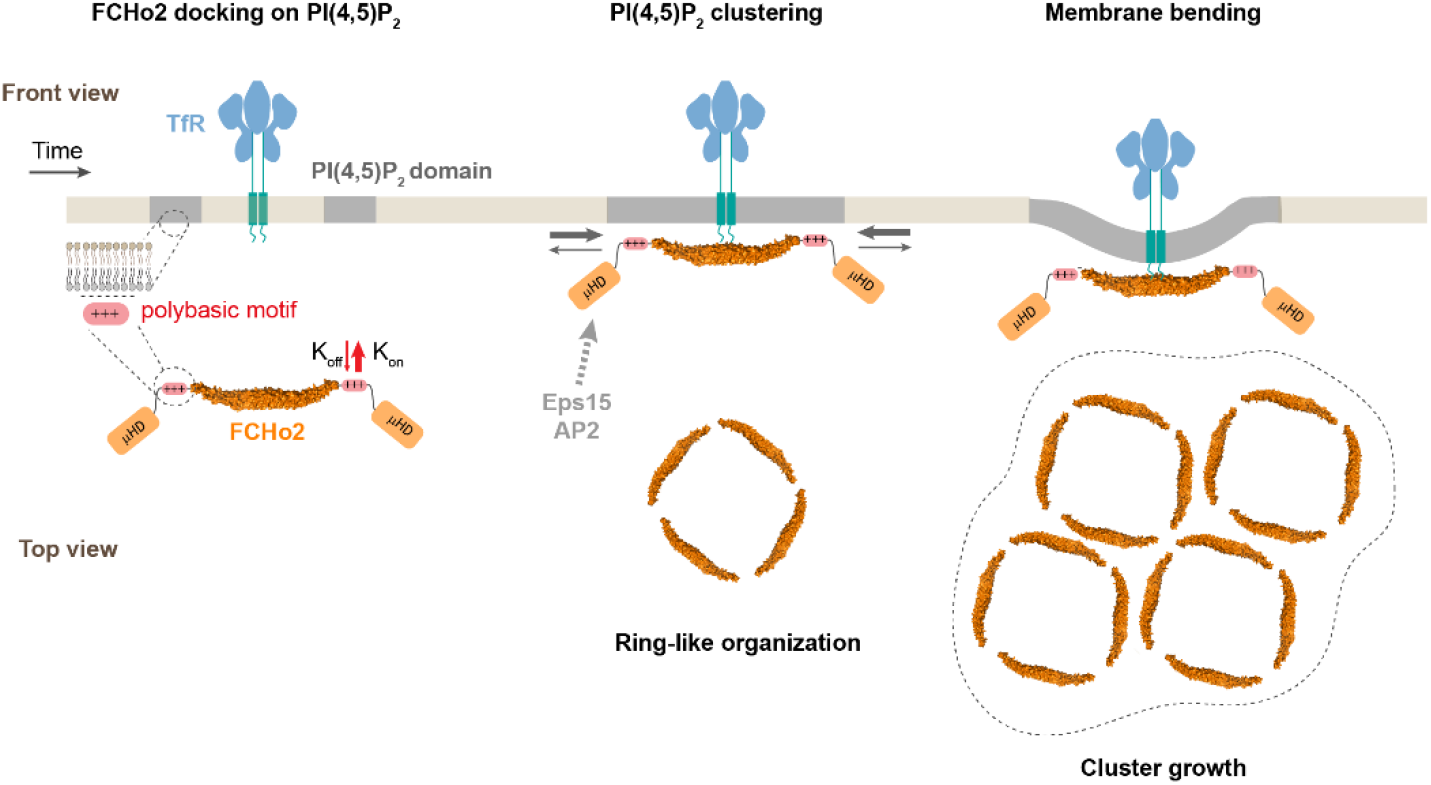
Model of FCHo2 docking and self-assembly on membranes. FCHo2 recruitment of membranes is mediated by PI(4,5)P_2_. Increase in the local PI(4,5)P_2_ concentration enhances the spatial accumulation of FCHo2 through multivalent interaction between the positively charged surface of the F-BAR domain and polybasic motif and PI(4,5)P_2_ molecules. Binding of FCHo2 on cellular membranes is convoyed by PI(4,5)P_2_ clustering formation at the boundary of clathrin-regulated cargo receptors and the self-assembly of FCHo2 into a ring-like shape protein complex. As a result of this singular organization, the local accumulation of PI(4,5)P_2_ and μHD of FCHo2 is likely to facilitate the formation of an interacting network with downstream partners, such as Eps15 and AP_2_(7, 8), ultimately leading to the assembly of clathrin structures on a PI(4,5)P_2_-rich interface that could be amplified by phosphoinositides metabolizing enzymes(3). In the absence of endocytic partners, FCHo2 rings grow into sub-micrometric molecular clusters leading to membrane bending and the partitioning on curved membranes.

This work provides the first evidence that FCHo2 self-assembles into ring-like molecular complexes on flat membranes. HS-AFM movies show that the ring formation process is relatively fast and takes place within less than 100 s, which might be compatible with the temporal scales reported during clathrin-coat assembly(1). This type of ring organization agrees with a side-lying conformation proposed for F-BAR scaffolds at low protein densities on flat surfaces(10) and the partitioning of FCHo1/2 at the edges of flat clathrin lattices(31). Lateral contacts stabilize the self-assembly of F-BAR domains on membrane tubules(10, 32). Our investigations point out that lateral interactions might occur on flat surfaces. We observed the anisotropic growth and formation of FCHo2 molecular clusters in the absence of other endocytic proteins. The formation of FCHo2 molecular complexes might confer structural stability to the ring-like organization and, importantly, show that F-BAR proteins can moderately bend flat membranes at high protein densities.

## Conclusions

In conclusion, our study provides a molecular picture of the recruitment and self-assembly of the early endocytic protein FCHo2 on membranes. While PI(4,5)P_2_-enriched domains act as docking sites to recruit FCHo2 on *in vitro* and cellular membrane, we found that the binding of FCHo2 promotes the local accumulation of PI(4,5)P_2_ at the vicinity of clathrin-regulated cargo receptors. As a result, in the absence of phosphoinositides-metabolizing enzymes, FCHo2 can enhance the formation of clathrin structures through a local PI(4,5)P_2_ enrichment, which could explain previous studies showing that FCHo1/2 depletion slows down the progression of cargo-loaded clathrin structures (8, 9, 12, 33). Because FCHo2 is among the early proteins recruited at endocytic sites, the discovery that FCHo2 self-assembles into rings convoyed by membrane bending and PI(4,5)P_2_ accumulation provides a fundamental understanding of the initiating mechanism of clathrin-mediated endocytosis(2).

## Materials and Methods

### Lipids and reagents

Natural and synthetic phospholipids, including POPC, POPS, Egg-PC, Brain-PS, Brain-PI(4,5)P_2_ and fluorescent TopFluor-TMR-PtdIns(4,5)P_2_ are from Avanti Polar Lipids, Inc. Oregon green 488-DHPE and Alexa Fluor 647 Maleimide labelling kit are from Invitrogen. Atto647N-DOPE and polyclonal rabbit anti-clathrin heavy chain (dilution 1:100; Cat. Nr. HPA059143) was from Sigma. Monoclonal rabbit anti-EGFR (dilution 1:1000; Cat. Nr. 4267) and polyclonal rabbit anti-phospho-EGFR (Tyr1045) (dilution 1:1000; Cat. Nr. 2237) were from Cell Signaling.

### EGFR-GFP and TfR-GFP stable cell lines

pRRL.sin.cPPT.SFFV-EGFP/IRES-puro was kindly provided by C. Goujon (IRIM, CNRS UMR9004, Montpellier, France), the EGFR-GFP vector was a gift from Alexander Sorkin, (Addgene plasmid #32751) and the pBa.TfR.GFP vector was a gift from Gary Banker & Marvin Bentley (Addgene plasmid # 4506). The GFP was replaced by a GFP-delta-ATG using these primers: 5’-gtatatatatGGATCCGTGAGCAAGGGCGAGGAG-3’ and 5’-CTCACATTGCCAAAAGACG-3’. GFP-delta-ATG fragment replaced the GFP- fragment in pRRL.sin.cPPT.SFFV-EGFP/IRES-puro using a BamHI-XhoI digestion. EGFR was amplified with these primers: 5’-caaatatttgcggccgcATGCGACCCTCCGGGACG-3’ and 5’-gtataccggttgaacctccgccTGCTCCAATAAATTCACTGCTTTGTGG-3’ and cloned into pRRL.sin.cPPT.SFFV-EGFP-delta-ATG/IRES-puro using NotI-AgeI to generate a fused EGFR-GFP protein.

Lentiviral vector stocks were obtained by polyethylenimine (PEI) -mediated multiple transfection of 293T cells in 6-well plates with vectors expressing Gag-Pol (8.91), the mini-viral genome (pRRL.sin.cPPT.SFFV-EGFR-GFP/IRES-puro) and the Env glycoprotein of VSV (pMD.G) at a ratio of 1:1:0.5. The culture medium was changed 6 hr post-transfection and lentivectors containing supernatants harvested 48 hr later, filtered and stored at −80°C.

The mini-viral genome (pRRL.sin.cPPT.SFFV-TFR-GFP/IRES-puro) and the corresponding lentivectors were generated as detailed in the case of the EGFR-GFP.

To generate a stable cell line HT1080 expressing EGFR-GFP or TfR-GFP, HT1080 cells (a gift from N. Arhel, IRIM, CNRS UMR9004, Montpellier, France) were transduced in 6-well plates using the supernatant of one 6-well plates of the lentiviral stock production detailed above. The culture medium was changed 6 hr post-transduction. Puromycin (1μg/ml) was added 48h after transduction. The percentage of GFP-expressing cells was enumerated by flow cytometry 72 hr after selection under puromycin.

HT1080 cells constitutively expressing the EGFR-GFP or TfR-GFP were cultured in DMEM GlutaMAX supplemented with 10% fetal calf serum, 100 U·mL^−1^ of penicillin and streptomycin and 1μg·ml^−1^ of puromycin at 37°C in 5%CO_2_. Cell lines were tested negative for mycoplasma.

### Protein purification and protein labeling

pGEX-6P-1 vector coding for the mouse full-length FCHo2 (aa 1-809) and F-BAR domain (aa 1-262) and human FBP17 (aa 1-592) were obtained from H.T McMahon (MRC Laboratory of Molecular Biology, Cambridge, UK). Proteins were subcloned into a pET28a vector with a PreScission protease cleaving site. Proteins were expressed in Rosetta 2 bacteria and purified by affinity chromatography using a HiTrap™ chelating column (GE Healthcare) according to the manufacturer’s instructions in 50 mM Tris at pH 8.0, 100 mM NaCl. Proteins were expressed overnight at 18°C using 1 mM IPTG. Proteins were then dialyzed overnight in a Slide-A-Lyzer dialysis cassette (MWCO 10,000) before Alexa Fluor 647 maleimide labelling following the protocol described by the manufacturer (Invitrogen). Protein concentrations were measured using a Bradford assay (Biorad).

Recombinant GST-eGFP-PH-domain (PLCδ1) detecting PI(4,5)P_2_ was purified as described in(25).

### Xenopus *laevis* egg extracts

Laid eggs are rinsed twice in XB Buffer (100mM KCl, 1mM MgCl2, 0.1mM CaCl2, 50mM sucrose and 10mM HEPES at pH 7.7) and subsequently dejellied with 2% cysteine solution pH7.8. Once dejellied they are extensively rinsed with XB buffer to completely eliminate cysteine solution.

Eggs are then recovered from a Petri dish and treated with Ca2+ Ionophore (final concentration 2 μg/ml) and 35 minutes later, they are crushed by centrifugation for 20min at 10,000g at 4°C. The cytoplasmic layer is collected and supplemented with Cytochalasin B (50ug/ml) Aprotinin (5ug/ml), Leupeptin (5ug/ml) and 10 mM Creatin Phosphate. Cytoplasmic extract is centrifuged again for 20 min at 10,000g. Extract are frozen and then used as described in Figure 2.

The energy mix consisted of 1.5 mM ATP, 0.15 mM GTPγS, 16.7 mM creatine phosphate and creatine phosphokinase 16.7 U·ml^−1^, as previously reported(34)

### Supported lipid bilayers

Lipid mixtures consisted of: 80-85% Egg-PC, 10-15% Brain-PS and 5-10% of Brain-PI(4,5)P_2_. The amount of total negatively charged lipids was kept to 20% for any of the mixtures containing phosphoinositides at the expenses of Brain-PS. If needed, fluorescent lipids were added to 0.2%.

For fluorescence microscopy experiments, supported lipid bilayers were prepared as described in(35). Experiments were performed by injecting 15 μL of buffer (20 mM Tris, pH 7.4, 150 mM NaCl and 0.5 mg·ml^−1^ of casein). Supported lipid bilayers were imaged on a Zeiss LSM880 confocal microscope.

For HS-AFM experiments, supported lipid bilayers were prepared following the method described in(36). Briefly, large unilamellar vesicles (LUVs, diameter ~ 100 nm) were obtained by extrusion of multilamellar vesicles of 85% POPC, 10% POPS and 5% Brain-PI(4,5)P_2_ in 20 mM Hepes, pH 7.4, 150 mM NaCl. LUVs were supplemented with 20 mM of CaCl_2_ and deposited onto freshly cleaved mica disks. Samples were incubated for 20 min at 60 °C and extensively rinsed with 20 mM Hepes, pH 7.4, 150 mM NaCl, 20 mM EDTA. Finally, bilayers were rinsed and keep under the imaging buffer, 20 mM Hepes, pH 7.4, 150 mM NaCl.

### Plasma membrane sheets

Unroofing of HT1080 cells stably expressing the EGFR-GFP was performed by tip sonication as reported in(37). Cells were rinsed three times in cold Ringer buffer supplemented with Ca^2+^ (155mM NaCl, 3 mM KCl, 3mM NaH2PO_4_, 5mM HEPES, 10mM glucose, 2 mM CaCl_2_, 1mM MgCl_2_, pH 7.2), then immersed 10 s in Ca^2+^-free Ringer buffer containing 0.5 mg·mL^−1^ poly-L-lysine. Cells were unroofed by scanning the coverslip with the tip sonicator at 10% of power under HKMgE buffer consisting of 70 mM KCl, 30mM HEPES, 5 mM MgCl_2_, 3mM EGTA, pH 7.2. Unroofed cells were kept in HKMgE buffer. Fluorescent labeling of plasma membrane sheets was performed immediately after unroofing by incubating the sample with 100 nmol of TopFluor-TMR-PtdIns(4,5)P_2_ suspended in 0.2% of absolute ethanol during 5 min, as reported in(38). Then, samples were extensively rinsed with HKMgE buffer and immediately imaged under the Zeiss LSM880 confocal microscope. Before addition of 1 μM of FCHo2-Alexa647, unroofed cells were rinsed with HKMgE buffer supplemented with 0.5 mg·ml^−1^ of casein.

### Immunofluorescence

Supported lipid bilayers were fixed in 3.7% PFA in PBS for 2 min at room temperature, then rinsed in PBS twice. Samples were stained for the primary antibody for 45 min at room temperature in 1% BSA. Then, the secondary antibody was incubated for 45 min. Finally, samples were extensively rinsed in PBS, then in sterile water and mounted with a Mowiol^®^ 4-88 mounting medium (Polysciences, Inc.). Montage was allowed to solidify in the dark for 48h before microscope acquisitions.

### Silica thin film nanostructuration

SiO_2_ vertical nanostructures were prepared on conventional borosilicate coverslips with precision thickness No. 1.5 (0.170 ± 0.005 mm), as previously reported(25, 39). Briefly, Si masters were elaborated using LIL Lithography as detailed in(39, 40). PDMS (polydimethylsiloxane) reactants (90 w% RTV141A; 10 w% RTV141B from BLUESIL) were transferred onto the master and dried at 70 °C for 1 h before unmolding.

Silica precursor solution was prepared by adding 4.22 g tetraethyl orthosilicate (TEOS) into 23.26 g absolute ethanol, then 1.5 g HCl (37%), and stirring the solution for 18 h. The final molar composition was TEOS:HCl:EtOH=1:0.7:25. All the chemicals were from Sigma. Gel films were obtained by dip-coating the coverslips with a ND-DC300 dip-coater (Nadetech Innovations) equipped with an EBC10 Miniclima device to control the surrounding temperature and relative humidity to 20°C and 45-50%, respectively. The thickness of film was controlled by the withdrawal rate at 300 mm/min. After dip-coating, gel films were consolidated at 430°C for 5 min. Then, a new layer of the same solution was deposited under the same conditions for printing with the PDMS mold. After imprinting, the samples were transferred to a 70 °C oven for 1 min and then to a 140 °C for 2 min to consolidate the xerogel films before peeling off the PDMS mold. Finally, the sol–gel replicas were annealed at 430 °C for 10 min for consolidation.

### Fluorescence microscopy

Images were acquired on a Zeiss LSM880 Airyscan confocal microscope (MRI facility, Montpellier). Excitation sources used were: 405 nm diode laser, an Argon laser for 488 nm and 514 nm and a Helium/Neon laser for 633 nm. Acquisitions were performed on a 63x/1.4 objective. Multidimensional acquisitions were acquired via an Airyscan detector (32-channel GaAsP photomultiplier tube (PMT) array detector). 3D images were acquired by fixing a 0.15 μm z-step in order to cover the entire surface of vertical nanostructures.

### HS-AFM imaging

HS-AFM movies were acquired with an HS-AFM (SS-NEX, Research Institute of Biomolecule Metrology, Tsukuba, Japan) equipped with a superluminescent diode (wavelength, 750 nm; EXS 7505-B001, Exalos, Schlieren, Switzerland) and a digital high-speed lock-in Amplifier (Hinstra, Transcommers, Budapest, Hungary)(41) following the protocol detailed at(42). Scanning was performed using USC-1.2 cantilevers featuring an electron beam deposition tip (NanoWorld, Neuchâtel, Switzerland) with a nominal spring constant k = 0.15 N/m, resonance frequency f(r) = 600 kHz, and quality factor Qc ≈ 2 under liquid conditions. For high-resolution imaging, the electron beam deposition tip was sharpened by helium plasma etching using a plasma cleaner (Diener Electronic, Ebhausen, Germany). Images were acquired in amplitude modulation mode at the minimal possible applied force that enables good quality of imaging under optical feedback parameters.

### Image processing and quantification

Line profiles of the fluorescence intensities were done using ImageJ(43) and the kymographs were made using the Kymograph plugin (http://www.embl.de/eamnet/html/body_kymograph.html).

Protein binding was quantified by measuring the mean grey value of the protein channel that was then normalized by the mean gray value of the membrane intensity (as indicated by the TF-TMR-PI(4,5)P_2_ fluorescence) in the same image. Mean gray values were measured once protein binding reached the steady-state, which was estimated from the binding kinetics to be < 4 min. Protein binding was averaged from 3 experimental replicates. Mean gray values were measured using ImageJ. Concentrations and confocal parameters were kept constant between experiments and samples.

The automatic analysis of images to determine molecular clusters was performed with ImageJ(43). Spots of different sizes are detected using a scale space spot detection (44) and overlapping spots are merged. The LoG filter of the FeatureJ plugin(45) is used to create the scale space. Starting points are detected as local minima of the minimum projection and as minima on the smallest scale. A simplified linking scheme is applied that looks for minima along the scales for each starting point within the radius of the spot on the given scale. Two spots are merged if at least 20 percent of the surface of one spot is covered by the other. The noise tolerance for the spot detection is determined manually for each series of input images.

To determine the height of each protein surface on SiO_2_ vertical nanostructures, we applied an automated image analysis procedure that calculates the statistics about the average maximal height of the signal in each channel of interest, as previously described(25). The analysis has been implemented as an ImageJ(43) macro toolset(46).

HS-AFM images were processed using Gwyddion, an open-source software for SPM data analysis, and WSxM(47).

### Data representation and statistical analysis

Representation of cross-section analysis and recruitment curves was performed using Origin software.

3D rendering of Airyscan images was generated with the 3/4D visualization and analysis software Imaris (Oxford Instruments).

Statistical analysis was performed using Prism GraphPad software.

## Acknowledgements

The authors thank H.T. McMahon for kindly providing F-BAR domain protein constructs. C. Goujon and O. Moncorgé for helping in the transduction of HT1080 cells. C. Holuka for assistance with plasma membrane sheets. P. Maiuri for assistance in data analysis. J.B. Manneville for critical reading of the manuscript and discussion. S. Roche, C. Favard, and D. Muriaux for scientific discussions. We acknowledge the imaging facility MRI, member of the national infrastructure France-BioImaging infrastructure supported by the French National Research Agency (ANR-10-INBS-04, «Investments for the future»). L.P. acknowledges the ATIP-Avenir program (AO-2016) and ANR-18-CE13-0015-02 for financial support. A.C-G. acknowledges the financial support from the European Research Council (ERC) under the European Union’s Horizon 2020 research and innovation program (No.803004).

## Author contributions

F.E-A and L.P conceptualized the study. F.E-A. I.C., D.S-F, R.R., A.C-G and L.P. performed experiments and/or analyzed results. F.E-A. and J.V. generated the recombinant proteins used in this study. T.L. and A.C. produced the *X. laevis* egg extracts. S.V participated in the production of plasma membrane sheets. V.B. designed the automated analysis of images. C.A-A. generated the HT1080 stable cell lines. F.E-A and L.P wrote the manuscript. All authors contributed to the last version of the manuscript.

## Supporting information

**Figure S1.**
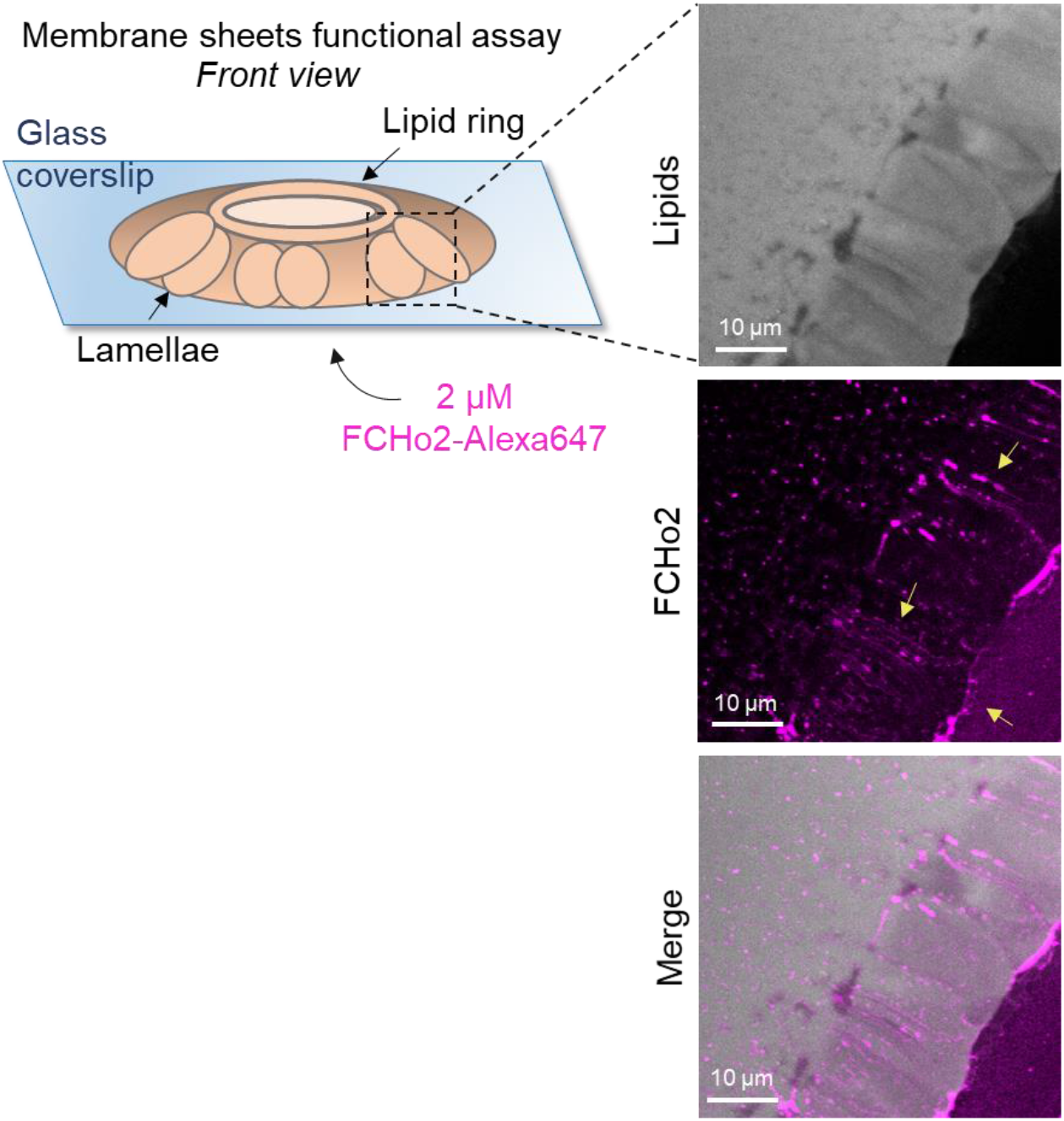
*Left*, schematic representation of the membrane sheets *in vitro* assay, as detailed by Itoh *et al*.(15), to test the functionality of recombinant full-length FCHo2-Alexa647. Membrane sheets were made using brain polar lipid extracts (Avanti). *Right*, representative still confocal images showing the formation of membrane tubes after addition of 2 μM of FCHo2, as indicated by yellow arrows.

**Figure S2.**
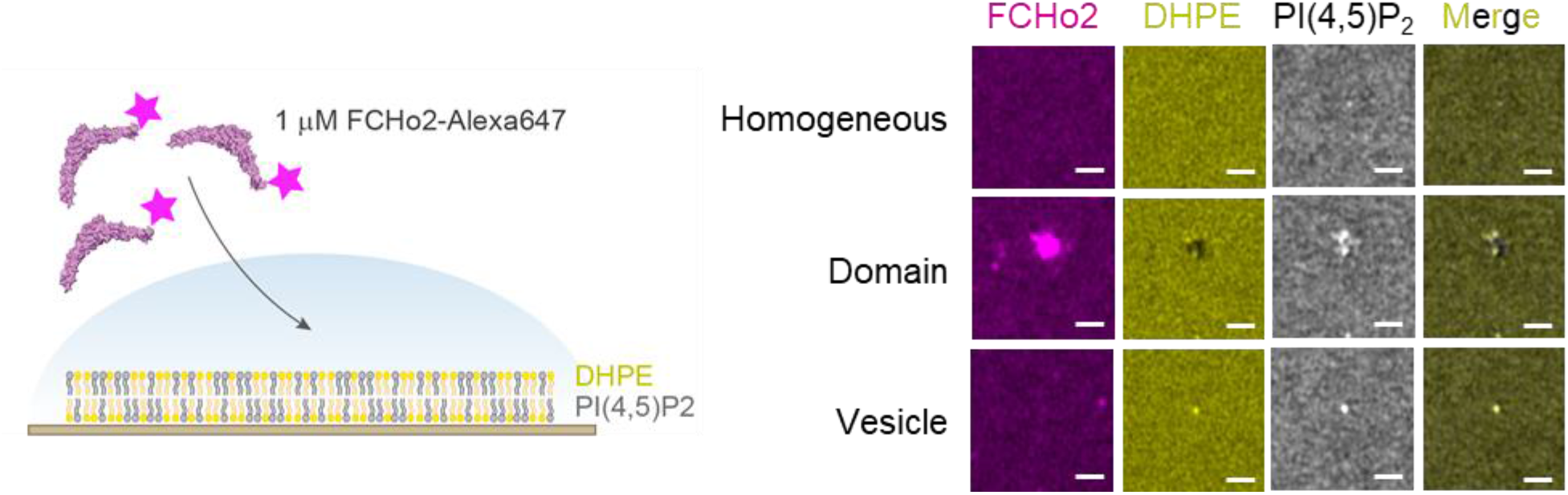
*Left*, cartoon of the experimental setup to detect the interaction of FCHo2-A647 with PI(4,5)P_2_-enriched domains. *Right,* representative airyscan images showing the lipid organizations that were detected under the experimental conditions: (i) an homogenous PI(4,5)P_2_ organization, (ii) the formation of lipid domains and (iii) the presence of adsorbed lipid vesicles. Scale bar, 1 μm.

**Figure S3.**
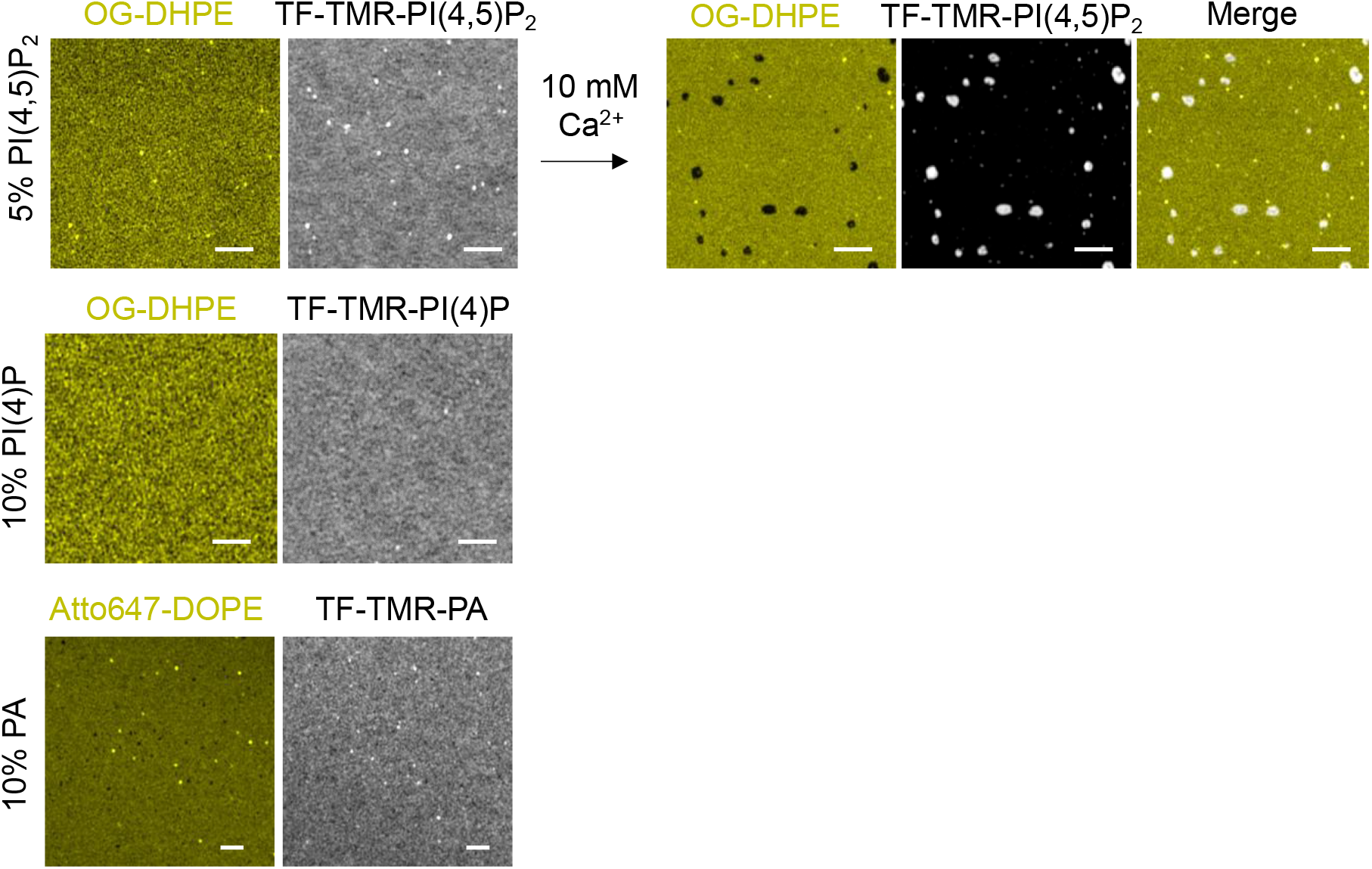
Representative airyscan images showing the lipid organization of different acidic TF-TMR lipid dyes (PI(4,5)P_2_, PI(4)P and phosphatidic acid (PA) in gray) on lipid bilayers containing 20% mol of total negative charge relative to a neutral lipid dye (OG-DHPE or Atto647-DOPE). Lipid dyes were added to a final 0.2% mol at the expenses of the non-labeled counterpart. Addition of 10 mM of Ca^2+^ on lipid bilayers made of 5% PI(4,5)P_2_ and doped with OG-DHPE and TF-TMR-PI(4,5)P_2_ shows the detection of PI(4,5)P_2_-enriched domains on this supported lipid bilayers. Scale bar, 2 μm.

**Figure S4.**
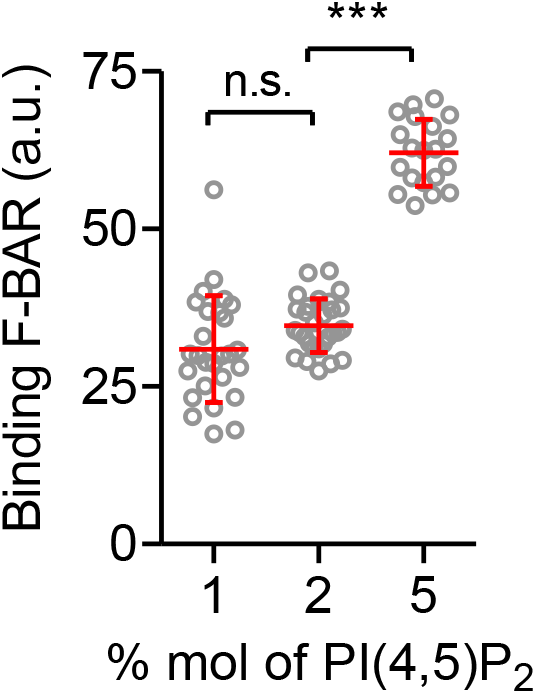
Binding of F-BAR on membranes containing different % mol of PI(4,5)P_2_. Mean ± s.d. is displayed in red and one-way ANOVA in black. n=27, n=29, and n=20 for 1%, 2%, and 5% mol of PI(4,5)P_2_ respectively.

**Figure S5.**
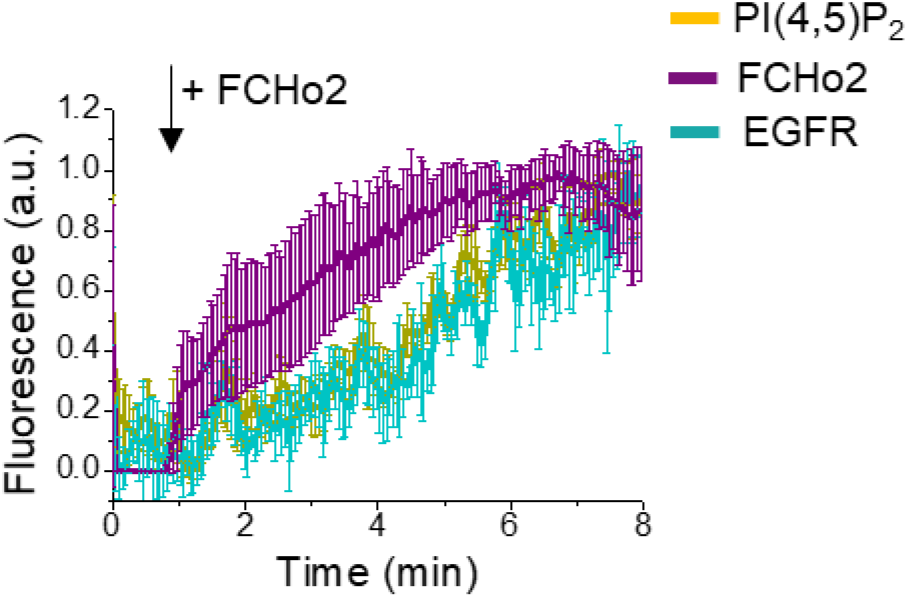
Fluorescence quantification over time of the EGFR (cyan), PI(4,5)P_2_ (yellow) upon injection of 1μM of FCHo2-Alexa647 (magenta) on EGFR-positive spots on plasma membrane sheets. Each curve represents the mean ± s.d., n = 7.

**Figure S6.**
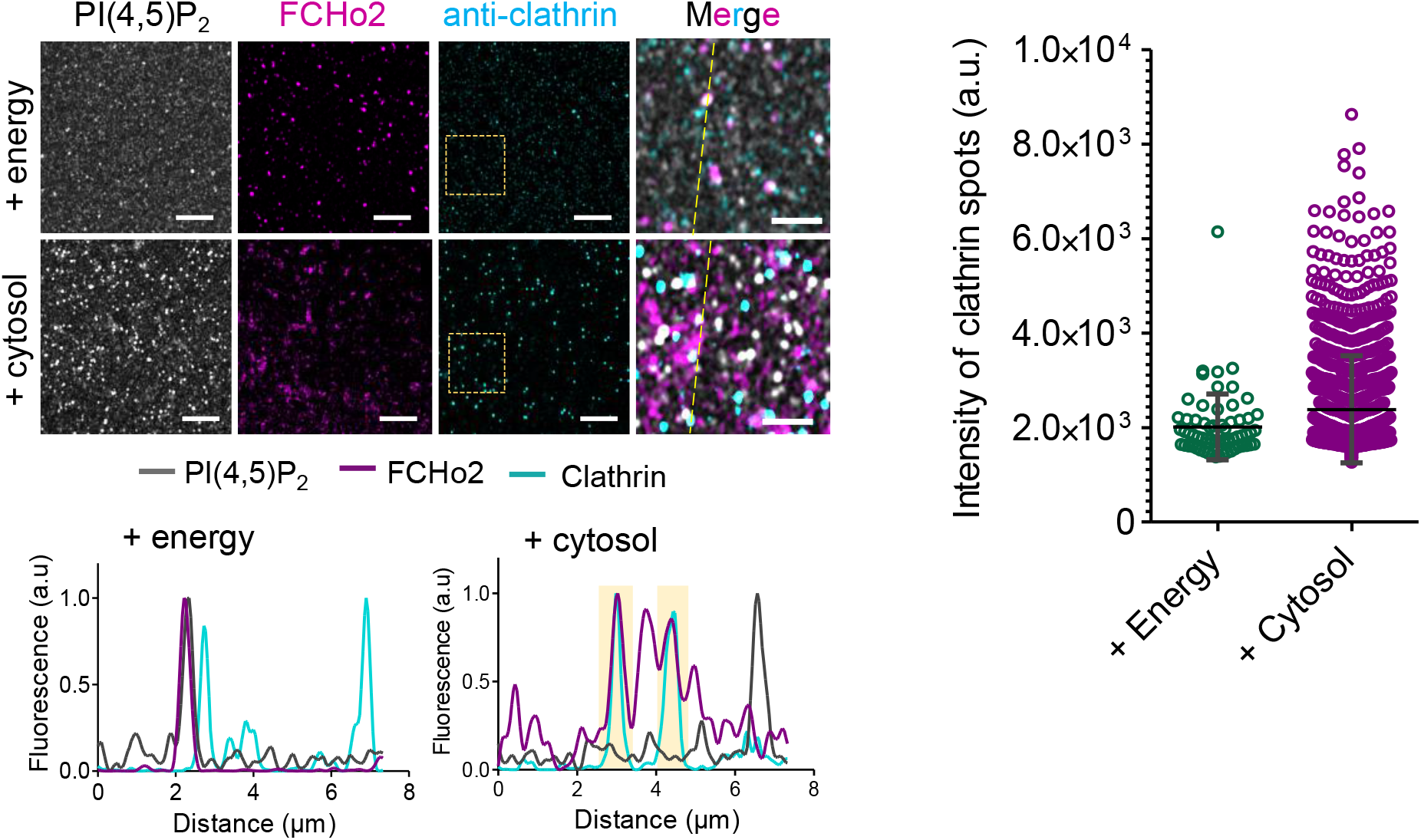
Representative airyscan images of the immunofluorescence assay showing the localization of PI(4,5)P_2_ (gray), FCHo2 (magenta) and clathrin (cyan, anti-clathrin antibody) on 5% PI(4,5)P_2_-containing lipid bilayers incubated with the energy mix alone (+ energy) or with the cytosolic extract and energy mix (+ cytosol). Sale bar, 5 μm.

